# BGP: Branched Gaussian processes for identifying gene-specific branching dynamics in single cell data

**DOI:** 10.1101/166868

**Authors:** Alexis Boukouvalas, James Hensman, Magnus Rattray

**Affiliations:** Division of Informatics, Imaging and Data Sciences Faculty of Biology, Medicine and Health University of Manchester; prowler.io

## Abstract

High-throughput single-cell gene expression experiments can be used to uncover branching dynamics in cell populations undergoing differentiation through use of pseudotime methods. We develop the branching Gaussian process (BGP), a non-parametric model that is able to identify branching dynamics for individual genes and provides an estimate of branching times for each gene with an associated credible region. We demonstrate the effectiveness of our method on both synthetic data and a published single-cell gene expression hematopoiesis study. The method requires prior information about pseudotime and global cellular branching for each cell but the probabilistic nature of the method means that it is robust to errors in these global branch labels and can be used to discover early branching genes which diverge before the inferred global cell branching. The code is open-source and available at https://github.com/ManchesterBioinference/BranchedGP.

## Background

Single-cell gene expression data can be used to uncover cellular progression through different states of a temporal transformation, e.g. during development, differentiation or disease. As single cell protocols improve, a flurry of methods have been proposed to model branching of cellular trajectories to alternative cell fates (Haghverdi *et al*., 2016; Setty *et al*., 2016; Qiu *et al*., 2017; Street *et al*., 2017). In these and similar methods, pseudotime is estimated and a global branching structure is inferred. Our focus in this paper is to propose a downstream analysis method that can sub sequently be used to model branching gene expression dynamics for individual genes. We are interested in discovering which genes follow the global cellular branching dynamics and whether these genes branch early or late with respect to the global cellular branching time. Recently, Qiu *et al*. proposed the branch expression analysis modelling (BEAM) approach that uses penalised splines to infer the individual gene branching time Qiu *et al*. (2017). Here we propose an alternative non-parametric method to model gene expression branching dynamics. We develop a probabilistic generative model of branching dynamics which can be used to assess the evidence for branching and provides a posterior estimate of the branching time. The posterior distribution over branching time can be used to identify the most likely branching time for each gene as well as an associated credible region capturing our uncertainty in the estimate.

Our approach is based on Gaussian processes (GPs) which are a class of flexible non-parametric probabilistic models. GPs have a long history in temporal and spatial statistics and have gained popularity in many areas of machine learning, including multivariate regression, classification and dimensionality reduction (Rasmussen and Williams, 2006). GPs have been used for dimensionality reduction of single-cell expression data (Buettner and Theis, 2012; Buettner *et al*., 2015) and more recently for pseudotime estimation where the effect of uncertainty in the inferred pseudotime can be quantified (Campbell and Yau, 2016) and capture time can be included as prior information (Reid and Wernisch, 2016). GP-based methods have also been used for modelling global cellular branching dynamics from single-cell data after assigning pseudotime to cells (Lönnberg *et al*., 2017).

Here, we build on the work of Yang *et al*. (2016) who developed a GP model for the identification of the time where two gene expression time course datasets first diverge from one another. They define a novel GP covariance function that constrains two functions to intersect at a single point. The divergence time is inferred by numerically approximating the posterior using a simple histogram approach. The model is used to identify when a gene first becomes differentially expressed in time course gene expression data under control and perturbed conditions. In their approach all data points have been labelled with the branch that generated them and the ordering of time points is assumed known. Although similar to the problem of modelling branching in single-cell data after pseudotime is inferred, this two-sample time series method cannot be applied directly to our problem because we have to allow for uncertainty in which branch each cell belongs to.

Also closely related to the present work, the overlapping mixture of GPs (OMGP) (Lázaro-Gredilla *et al*., 2012) is a mixture model for time-series data where the mixture components are GP functions and data at any time can be assigned to any of the components. In the case of single-cell data, after pseudotime is assigned to each cell then the OMGP model can be used to assign cells to different trajectories. The cell labels do not have to be known in advance and can be inferred through fitting the model to data. However, the OMGP models the cellular trajectories as independent rather than branching. The OMGP model has been applied to single cell data to infer global cell branching times (Lönnberg *et al*., 2017) but as the OMGP assumes the latent functions are independent without any branching, a heuristic based on a piecewise linear fit of the log likelihood surface is proposed to identify the most likely branching times. This is problematic since the OMGP does not provide a proper generative model of branching dynamics and therefore it is not clear how to compute the posterior distribution over the branching time.

Our main methodological contribution here is to generalise the OMGP model to explicitly account for dependence between the functions in the mixture model. Specifically, we consider the case where the functions branch as in Yang *et al*. (2016). This allows us to develop a probabilistic model over branching cellular trajectories where the assignment of cells to branches is not known in advance. Our new model allows us to calculate the posterior distribution over branching time for each gene while allowing for uncertainty in the branch labels for each cell. This uncertainty is especially important for early-branching genes, since cells are not labelled with a branch prior to the global cellular branching time which we assume is known.

A naive implementation of GP models scales cubically with the size of the data. As increasing numbers of cells can be profiled in new single cell protocols, we ensure the scalability of our approach by employing two complementary approaches. Firstly we use sparse inference (Quiñonero-Candela and Rasmussen, 2005; de Garis Matthews, 2016) that allows model fitting to scale with the number of inducing points. The latter is a user-defined value that trades off model accuracy and training time. Specifically for *N* cells, naive covariance inversion scales as *O*(*N*^3^) while under sparse inference with *k* inducing points it scales as *O*(*k*^2^*N*). Secondly, we provide an open-source implementation that leverages the GPflow library (Matthews *et al*., 2017), which both simplifies the implementation due to automatic symbolic differentiation and allows for the necessary matrix operations to be computed in parallel across many CPU nodes or GPUs.

## Results and discussion

### Overview of BGP

Before applying our algorithm we require the pseudotime for each cell and the global branching pattern of the cells to be established. For pseudotime estimation we use the reversed graph embedding approach Qiu *et al*. (2017), termed DDRTree, which we have found to be effective in recovering pseudotime in the presence of branching, although other approaches may be selected. Qiu *et al*. Qiu *et al*. (2017) have shown the reverse graph embedding approach to outperform DPT (Haghverdi *et al*., 2016), Wish-bone (Setty *et al*., 2016) and other methods in their analysis. The DDRTree method assigns each cell to a branch and identifies a set of globally defined branching points across all genes.

The overlapping mixture of Gaussian processes (OMGP) (Lázaro-Gredilla *et al*., 2012) is a mixture model of independent Gaussian processes (GPs) that is able to associate an observation with the generating GP. The authors term this the association problem and derive a variational inference algorithm for the case of independent GPs. Our work extends the OMGP model in two directions: firstly, we remove the assumption of latent function independence and allow dependent GPs as required by a branching model; secondly, we provide a sparse inducing point approximation that allows for scalable inference.

Let *F* be a branching GP evaluated for *N* data points with *M* branches and *Z ∈* {0, 1}*^N×M^* indicates which branch each cell comes from. The likelihood is *p* (*Y |F, Z*) = *𝒩 (Y |ZF, σ*^2^*I*) and as in Lázaro-Gredilla *et al*. (2012) we place a categorical prior on the indicator matrix 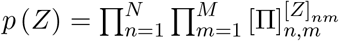 we place a GP prior on the latent functions *p* (*F | t_b_*) = *𝒢𝒫* (0, *K| t_b_*) which constrains the latent functions to branch at pseudotime *t_b_*. Note that the latter does not factorize as in Lázaro-Gredilla *et al*. (2012) as the latent functions are dependent. Full details on the model derivation and inference scheme used, including the inducing point approximation, is provided in the Methods section.

Global branching labels such as those provided by DDRTree can provide an informative prior *p*(*Z*) for all genes. The prior before the global branching point is uninformative as no global assignment is available. This is relevant for early-branching genes which may start branching earlier than the global cellular branching. After the global branching point, the prior favours increased assignment probability to the globally assigned branch. However, as the prior we use places non-zero mass on the alternative assignment, the resulting assignment may differ from the global allocation given enough evidence from the likelihood term. This allows the model to correct mislabelled cells as well as account for sources of noise in the data such as dropout for lowly expressed genes. The simplest construction of the prior, as used in this paper, is to specify a common uncertainty for all cells based on the global branching labels. For our results we have assigned a fixed probability for all cells of 80% of belonging to the globally assigned branch. However other constructions are possible; for instance the distance of the cells from the global branching time may be used to adjust the associated prior uncertainty.

The model hyperparameters are fitted by maximising a bound on the log-likelihood. The log-likelihood is not analytically tractable as it involves integrating out the indicator matrix *Z* and therefore we use a variational approximation. A lower bound is available using Jensen’s inequality

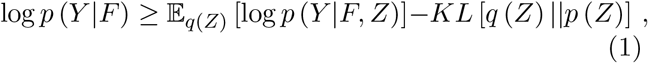

Where we use a mean-field approximation *q* (*Z, F*) = *q* (*Z*) *q* (*F*) with the latent functions *F* independent of the association indicators *Z* and *q*(*Z*) = ∏_*nm*_φ_*nm*_. The *φ_nm_* approximates the posterior probability of cell *n* belonging to branch *m*. The latter is either the trunk state or one of the two branches in the case of a single branching considered in the applications here. Then *F* can be integrated out to get the marginal likelihood *p*(*Y*).

The branching time posterior probability is calculated using the approximate marginal likelihood evaluated at a set of candidate branching points *S_B_* of size *N_b_*. The posterior for a candidate branching time *c* is

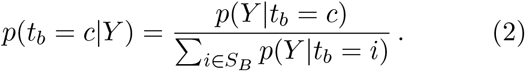

We can also calculate a likelihood ratio of branching versus not branching to rank genes by how likely their expression exhibits branching. By numerically integrating out the uncertainty of the branching location, the ratio statistic includes the effect of posterior uncertainty:

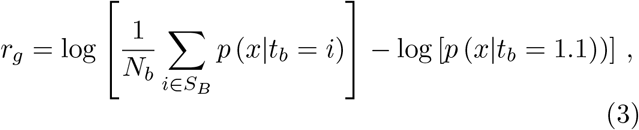

where *t_b_* = 1.1 specifies the model does not branch as the pseudotime is specified in [0, 1].

An example of the fit is shown in Figure 1 (b) where the uncertainty in the cell branch association is shown in conjunction with the posterior on the branching times. For visualisation the cell assignment to the top branch is shown. We see that most cells away from the branching point are assigned with high confidence to one of the branches. However, cells that are equidistant from both branches have high assignment uncertainty (0.5). This is also the case for cells close the branching location where the two branches are in close proximity. In the bottom panel of the figure, the posterior on the branching location is also shown. In this instance there are only two grid locations where the branching is likely to occur. This is reflected in Figure 1 (a) in the branching time uncertainty (magenta). The cell assignment uncertainty is incorporated in the branching time posterior; in cases where the branches separate quickly the posterior branching time uncertainty is likely to be small. This reflects one of the main benefits of employing a probabilistic model to identify branching dynamics as the assignment uncertainty is considered when calculating the branching time posterior. The cell assignment is inferred in the BGP model in contrast to model in Yang *et al*. (2016) where the assignment is assumed known.

**Figure 1:**
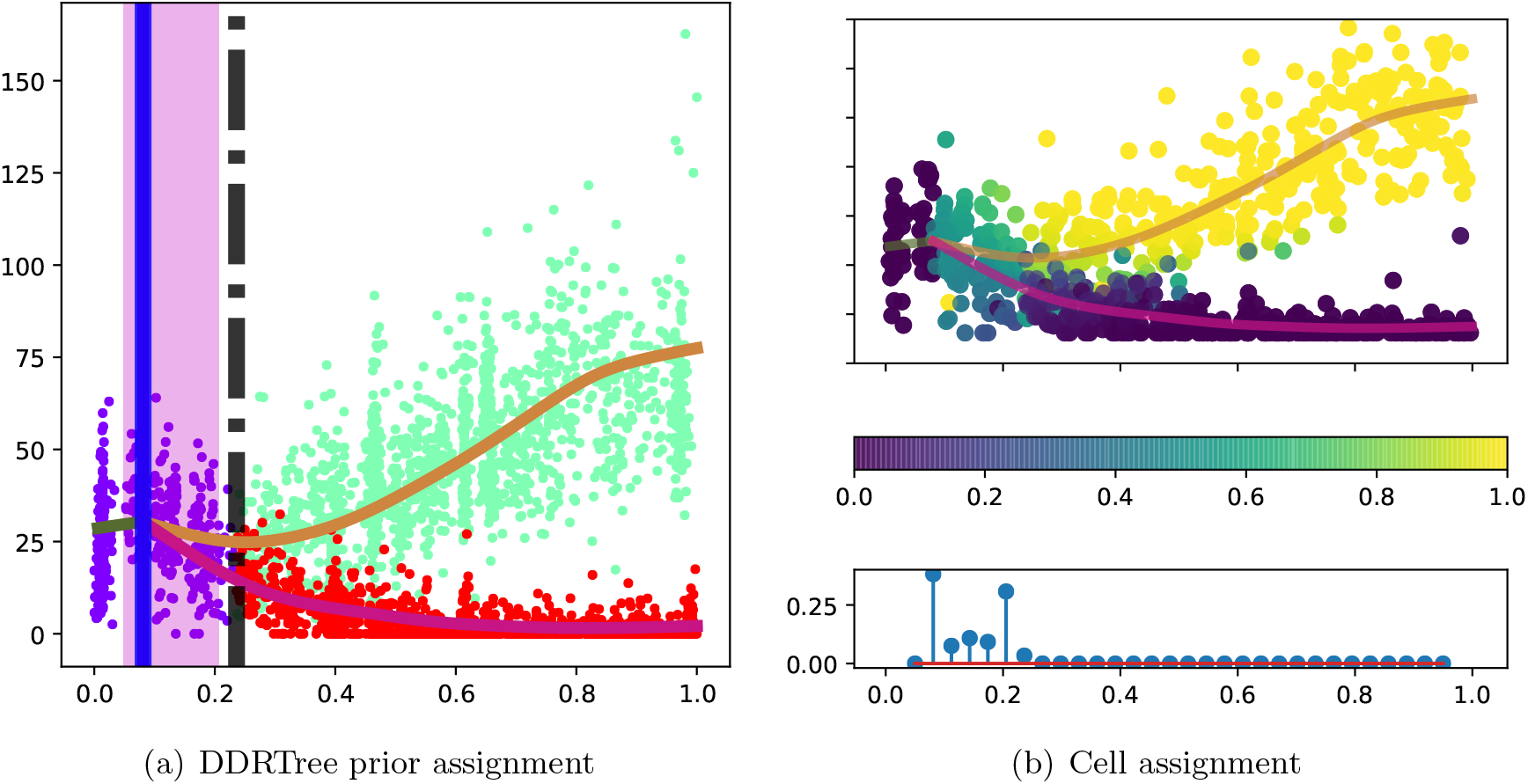
Hematopoiesis gene expression: BGP fit for MPO marker gene. In panel (a) the Monocle-DDRtree branching assignment is shown for each cell along with the global branching time (black dashed), most likely branching time (blue solid) and posterior branching time uncertainty (magenta background). In panel (b) the posterior cell assignment is shown in top subpanel. In the bottom subpanel the posterior branching time is shown.

### Synthetic study

We evaluate three methods, the mixture of factor analysers (MFA) (Campbell and Yau, 2017), the BEAM approach (Qiu et al., 2017) and the branching GP (BGP) model on synthetically generated data. For the synthetic study we use Gaussian noise and therefore we use the BEAM algorithm with a Gaussian likelihood function. MFA also assumes a Gaussian likelihood function. Data is generated from a branching Gaussian process with signal variance *σ*^2^ = 2, lengthscale *λ* = 1.2 and a range of noise levels (Table 2). Samples where the functions are crossing after the branching point were rejected since these may be difficult for other methods, e.g. BEAM identifies the last crossing location for its fitted splines and may therefore identify the wrong point in a time-series that crosses after branching. We address this issue in the real data study considered in the next section but do not consider it in the synthetic benchmark. We generate *N* = 150 data points with *D* = 40 genes and pseudotime in unit interval [0, 1]. The genes are separated in three groups depending on their branching behaviour and time (Table 1).

**Table 1:**
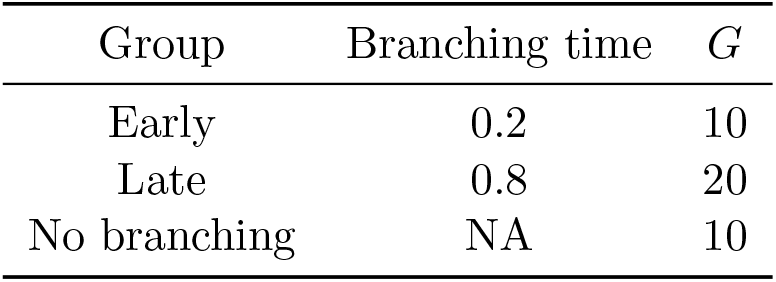
Synthetic gene groups. Branching times and number of genes *G* for each group. All scenarios use *N* = 150 cells and a total of *D* = 40 genes.

| Group | Branching time | $G$ |
| --- | --- | --- |
| Early | 0.2 | 10 |
| Late | 0.8 | 20 |
| No branching | NA | 10 |

**Table 2:** Synthetic study: Pseudotime rank correlation to the true time for both MFA and Monocle under both scenarios.

| Noise | MFA | Monocle |
| --- | --- | --- |
| 0.001 | -0.96 | 1.0 |
| 0.01 | 0.98 | 1.0 |
| 0.03 | -0.93 | 1.0 |
| 0.08 | -0.97 | 1.0 |
| 0.1 | -0.97 | 1.0 |
| 0.2 | 0.91 | 1.0 |

All methods were run with default parameter settings so it may be possible to improve on their performance by tuning these parameters; for example as in Campbell and Yau (2017) we found the performance of the algorithm dependent on the initialisation used. We contrast the performance of the BGP model both without a prior on cell assignment and an informative prior (80% prior probability) on cell assignment derived from the global Monocle assignment.

We first compare the pseudotime estimation accuracy of Monocle and MFA. Both methods achieve good performance as measured by the rank correlation of the estimated pseudotime to the ground truth (Table 2).

The log likelihood ratio of the branching GP can be used to rank the evidence of branching for each gene. Similar measures exist for the MFA and BEAM method. We first compare the three methods on their ability to discriminate branching from non-branching genes (Table 3). The metric we use is the area under the curve (AUC) which provides a reasonable measure when the number of positives and negatives in the ground truth are balanced. Both BEAM and BGP have higher accuracy than MFA whose performance varies significantly. The inclusion of an informative prior improves the performance of the BGP model resulting in consistently high performance for all noise levels. The performance of BEAM decreases with increased noise level which is also the case for the BGP model to a lesser extent.

**Table 3:**
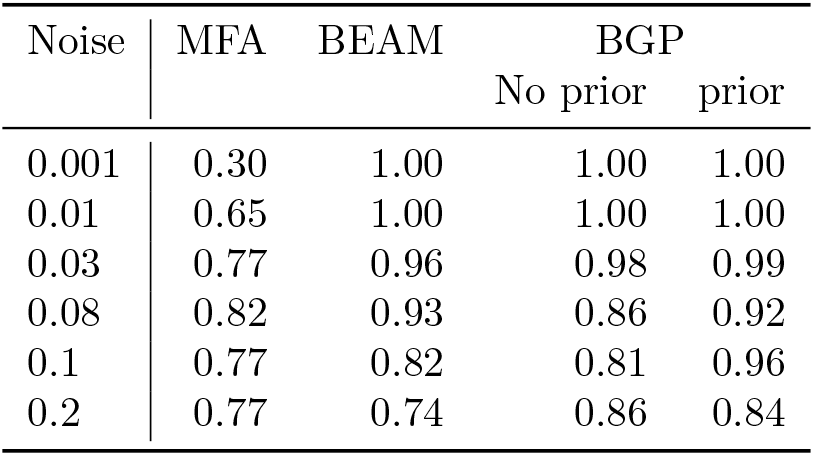
Synthetic study: Area under the curve (AUC) for detecting branching genes

We also examine the error in identifying the branching time. As MFA does not provide such an estimate, we only consider the BEAM and BGP methods. The error in estimating branching time for the BEAM and BGP methods is given in Table 4. The error for the BGP method is consistently lower than the BEAM method. The informative prior allows for more consistent performance of the BGP method with substantial increases in accuracy in some scenarios, especially so for the highest noise level (0.2) where the error is reduced from 0.15 to 0.08. The lack of robustness of the BEAM approach to high noise is demonstrated in Figure 2. In the low noise scenario (Figure 2 (a)-(b)), both BEAM and BGP are able to recover the gene expression branching dynamics. In the high noise scenario (Figure 2 (c)-(d)), the global branching time is early due to the presence of early branching genes in the data. The later branching gene depicted has a branching time of *b* = 0.8 and the global assignment correctly separates the two branches. However due to the high noise in the data the spline is unable to identify the correct branching time and significantly underestimates the branching time (Figure 2 (c)). In contrast the BGP model correctly identifies the late branching nature of the gene despite the early global branching time and the high noise level of the data (Figure 2 (d)).

**Table 4:** Synthetic study: Root mean squared error (RMSE) for branching time estimation. Only performed on branching genes

| Noise | BEAM | BGP |  |
| --- | --- | --- | --- |
|  |  | No prior | prior |
| 0.001 | 0.12 | 0.03 | 0.03 |
| 0.01 | 0.11 | 0.04 | 0.05 |
| 0.03 | 0.13 | 0.06 | 0.07 |
| 0.08 | 0.20 | 0.14 | 0.09 |
| 0.1 | 0.23 | 0.12 | 0.09 |
| 0.2 | 0.23 | 0.15 | 0.08 |

**Figure 2:**
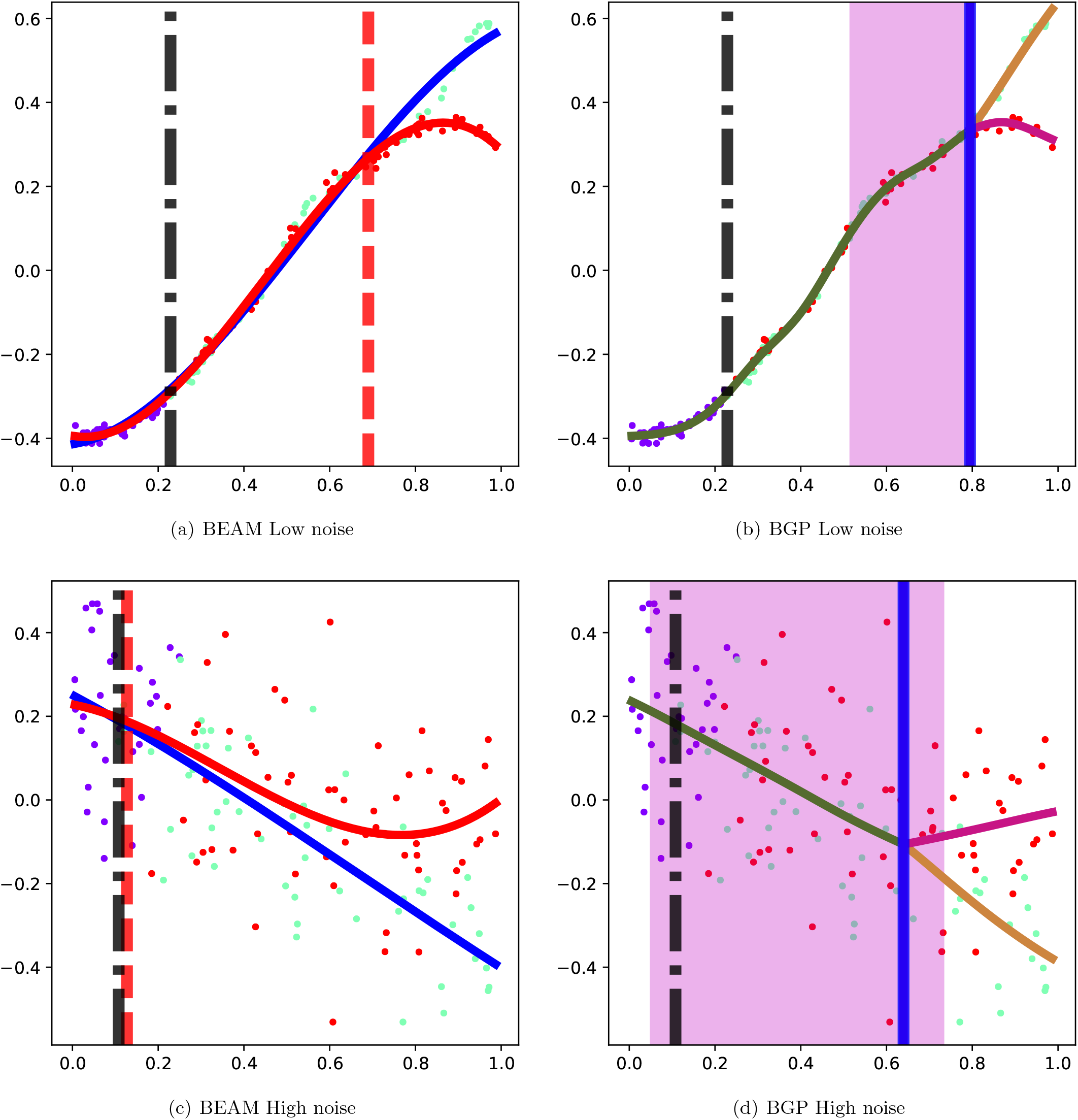
Synthetic data: Example BEAM and BGP model predictions for a late branching genes which branch at b=0.8. The vertical black bar is the global branching time. The vertical red bar is the BEAM branching time estimate, the vertical blue bar the BGP estimate and cells have been coloured by the global Monocle assignment.

More generally, the spline approach taken in BEAM suffers from a consistent bias in branching time estimation that pulls all estimates towards the global branching time. To clearly demonstrate this effect we examine an additional synthetic example with 3 genes branching very early (0.1), 27 genes branching late (0.7) and 10 genes not branching and select a low noise level (0.001).

.The late global branching time (Figure 3) due to the presence of a majority of late branching genes and few early branching genes, helps to clearly demonstrate the bias effect. As can be seen in the Figure 3 (a), the estimates for the BEAM method are biased towards the global branching time. The underestimation of branching times in BEAM for genes that branch later than the global branching time is most likely due to the spline regularisation employed by BEAM that tends to over-smooth the spline fit. The overestimation of branching times for early branching genes is due to the arbitrary assignment of cells prior the global branching time as no labels are provided by the global algorithm and no estimation is performed by the spline-fitting algorithm; see Figure 3 (c) for an illustrative example. The former could possibly be rectified by tuning the regularisation approach employed but the latter is a fundamental restriction of the BEAM approach that does not directly estimate branching assignments but only uses the globally derived label estimates. The BGP approach (Figure 3 (b)) does not suffer from this deficiency as the branch assignment is performed on a gene by gene basis at the cost of increased computation time. We note however the problem is parallelisable as each gene is treated independently.

**Figure 3:**
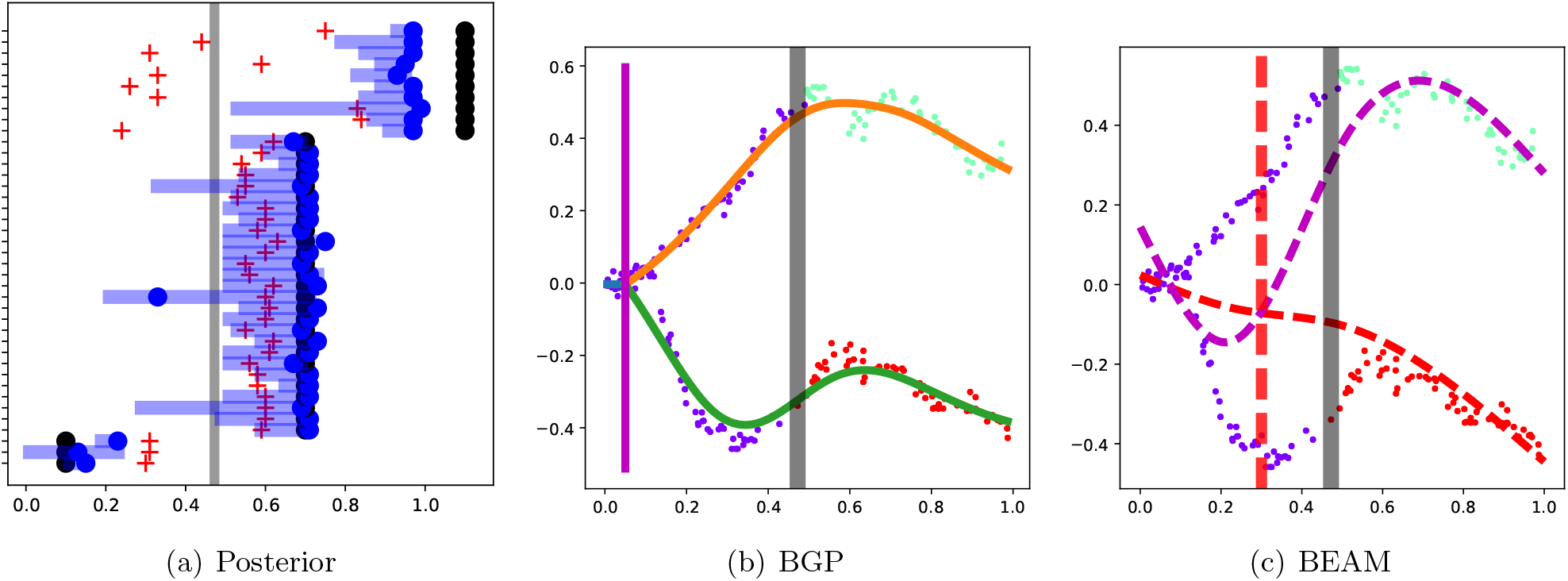
Synthetic data: fitting BGP and BEAM on an early branching gene. (a) The true branching times (black dots), BEAM times (red crosses) and BGP mean (blue dots) and 98% credible regions are shown. (b)-(c)The vertical grey bar is the global branching time. The vertical red dashed bar is the BEAM branching time estimate, the vertical magenta bar the BGP estimate and cells have been coloured by the global assignment.

Lastly, we examine the effect of poor state estimation on the BEAM and BGP methods (Figure 4). In Figures 4 (a)-(c) the Monocle state estimation accurately identifies the underlying branching dynamics and both BGP and BEAM correctly estimate the branching dynamics. In Figures 4 (d)-(f) we show an example of the effect of poor state estimation. The state estimation correctly identifies a single branching point but the majority of cells are assigned to one of the branches (red). As one of the global branches (red) spans both gene expression branches, it is unsurprising the spline approach fails to correctly identify the branch location and in fact overestimates the true branching time of *t_b_* = 0.2 (Figure 2 (a)). The corresponding BGP inference (Figure 2 (f)) overcomes the errors in global state estimation and the confidence interval includes the true branching time.

**Figure 4:**
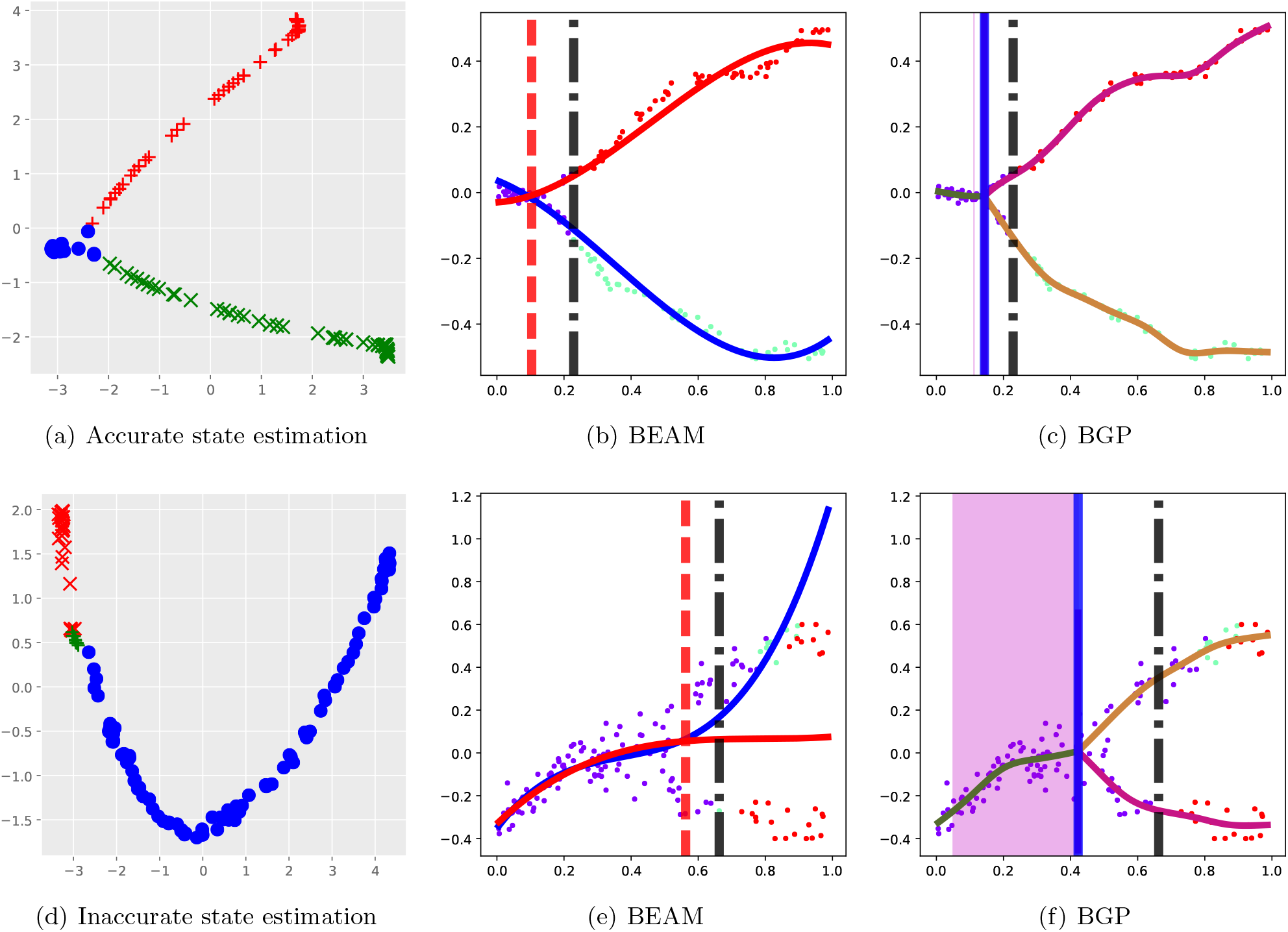
Synthetic data: Effect of Monocle global state estimation on BEAM and BGP model predictions for an early branching genes (b=0.2). Two different examples shown corresponding to accurate and inaccurate state estimation by Monocle. The vertical black bar is the global branching time. The vertical red bar is the BEAM branching time estimate, the vertical blue bar the BGP estimate and cells have been coloured by the global Monocle assignment.

The robustness of the BGP model can be understood in terms of the probabilistic nature of the model where both prior global state information is considered as well as a likelihood term that fits a branching process. Therefore the BGP prior model incorporates the best of both worlds; inclusion of global assignment information and assessment of cell assignment based on individual gene expression.

### Hematopoiesis single cell RNA-seq

We apply the BGP model on single-cell RNA-seq of hematopoietic stem cells (HSC) differentiating into myeloid and erythroid precursors (Paul *et al*., 2015). The data processing steps are described in the Methods section. The root state was selected using marker genes for common myeloid progenitors (CMP), megakaryocyte-erythroid progenitors (MEP) and granulocyte-macrophage progenitors (GMP). The two branches are clearly distinguishable in the latent space (Figure 5). The common myeloid progenitor FLT3 is highly expressed in the root of the tree whereas the MEP marker KLF1 and GMP marker MPO are expressed in each branch respectively. We also applied the Wishbone approach (Setty *et al*., 2016) and the two approaches have good agreement in the estimated pseudotime with a high rank correlation (0.92) and significant overlap in branching assignments.

A probabilistic model is an appropriate choice for early hematopoiesis which has been described as a cellular continuum of low-primed HSCs (Velten *et al*., 2017). The continuum contains transitory states rather than discrete progenitor cell types with some cell state transitions and lineage combinations more likely to occur than others. A probabilistic model such as BGP better reflects the probabilistic nature of lineage selection highlighted in Velten *et al*. (2017). In the BGP model in particular, each cell is associated with an allocation probability for each branch. The branching point can be interpreted as the earliest pseudotime from which probabilistic biases in lineage selection can be detected.

**Figure 5:**
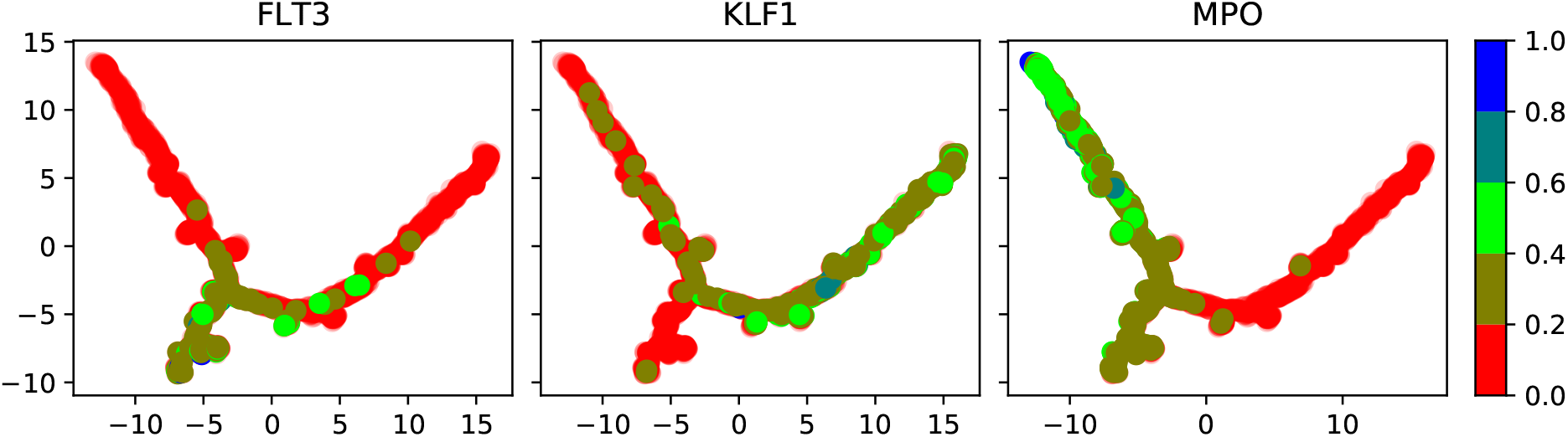
Hematopoiesis marker genes in Monocle latent space.

We find 737 genes out of a possible 1343 that show evidence of branching based on the log likelihood ratio statistic (Equation (3)). The posterior branching times for all branching are shown in Figure 6 (a). A significant portion of the spline branching times are near the end of pseudotime due to transitory gene expression which we discuss below. In Figures 6 (b)-(c) we show the branching times for ten marker genes that have been found to show significant evidence of branching. The colours reflect which branch is up-regulated after the most-likely branching time. For all GMP markers the same branch is upregulated (brown) whereas for the MEP markers the alternative branch is upregulated. Gene expression profiles are shown in Figure 1 for the MPO gene and Figure 7 for the other GMP markers. The MPO, PRTN3 and CTSG markers are highly expressed and show clear branching behaviour; as a result they are the three top-ranked genes in terms of the log likelihood ratio statistic (Equation (3)). In contrast, the CEBPA marker is lowly expressed and ranked 185th as the branches are less clearly separated. The profiles for some of the GMP markers are shown in Figure 8. When examining the individual gene expression for the APOE marker (Figures 8 (c)(d)), the gene expression exhibits a transitory phase where the magenta branch is initially upregulated after the branching event but then is downregulated. This behaviour turns out to be quite common and can lead the spline-based approach to fail (Figure 8 (d)). This is because after the transitory phase is completed, it is likely the spline predictions for each branch will intersect again. In the spline approach the latest intersection point between the two fitted splines is selected as the branching point (Qiu *et al*., 2017). This results in identifying the post-transitory phase point as the branching point. Transitory gene expression is also evident for other genes (Figure 9) wherein the spline approach similarly identifies the post-transitory phase as the dominant branching point.

**Figure 6:**
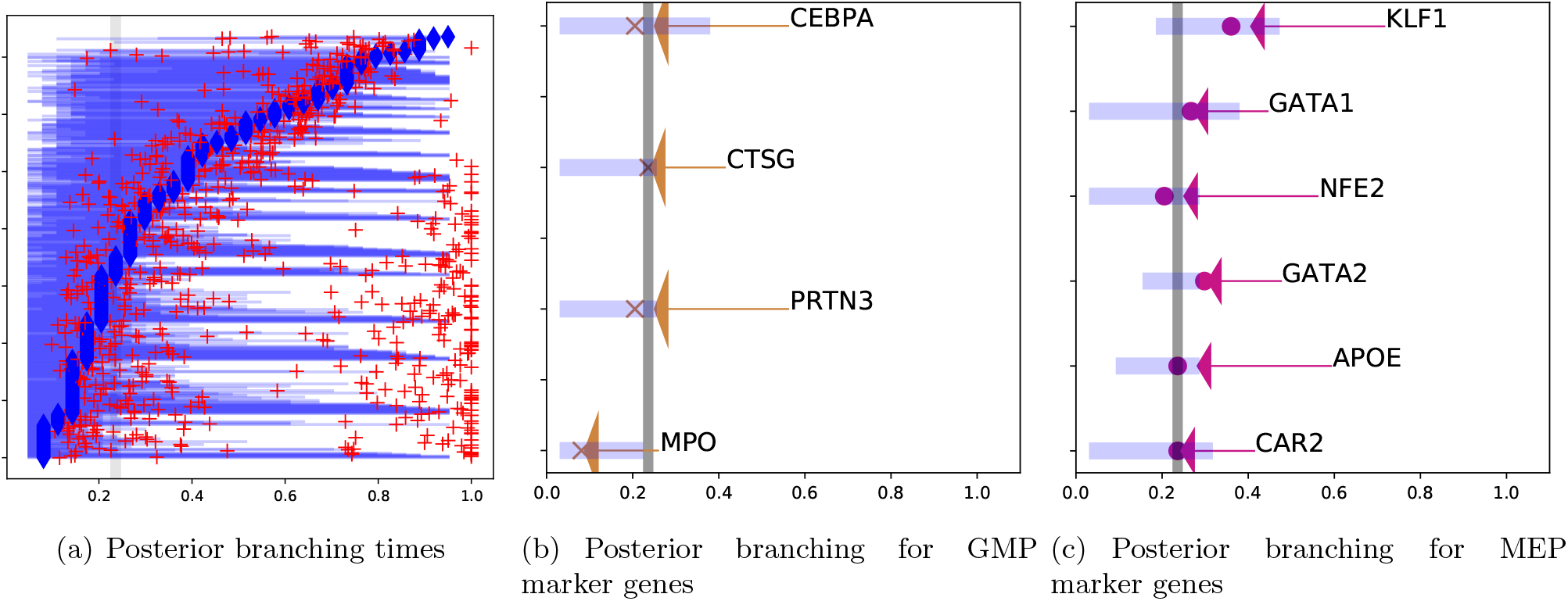
Posterior summary of 737 genes identified as branching and selected marker genes for the hematopoiesis data. The genes are ordered by the branching location. The spline estimation is shown as red crosses and the global branching time by a vertical grey bar (b=0.21).

**Figure 7:**
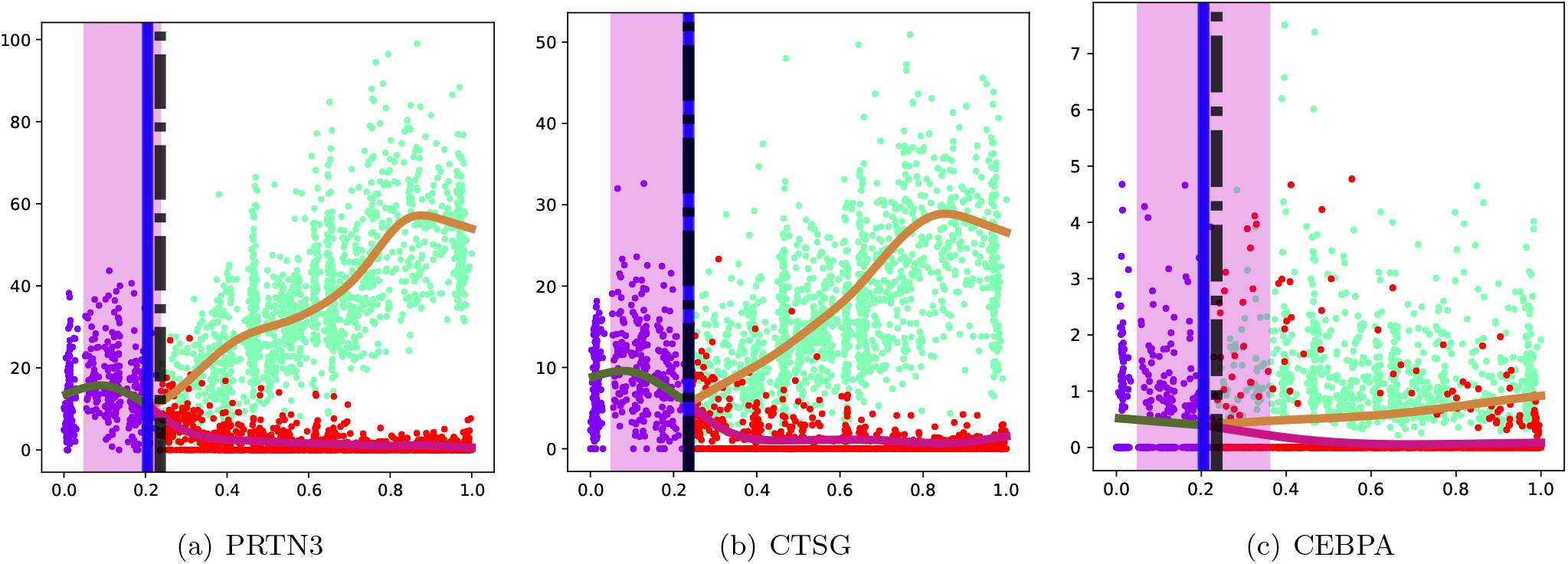
Hematopoiesis gene expression: BGP estimation for GMP marker genes. The global branching time (dashed black vertical line), BGP branching point mode (blue vertical line) and 98% posterior intervals (magenta vertical span) are also shown. The cells are marked according to the global allocation estimated by the DDRTree algorithm.

**Figure 8:**
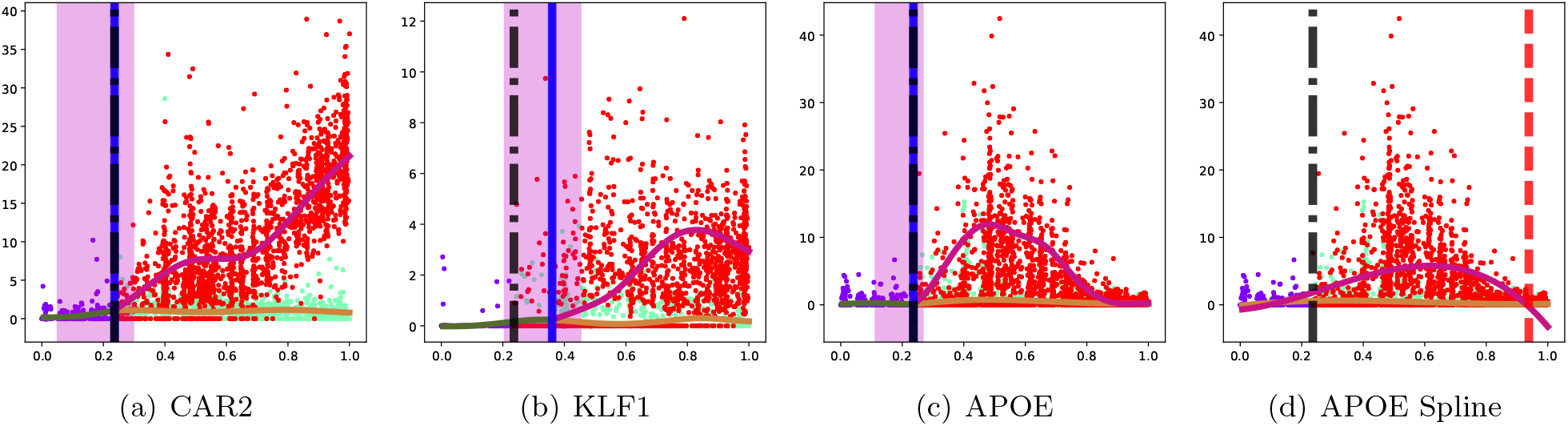
Hematopoiesis gene expression: BGP estimation for MEP marker genes. Also shown the spline fit for the APOE gene which exhibits transitory gene expression in one of the branches. The global branching time (dashed black vertical line), spline branching point mode (red vertical line) and 98% posterior intervals (magenta vertical span) are also shown. The cells are marked according to the global allocation estimated by the DDRTree algorithm.

**Figure 9:**
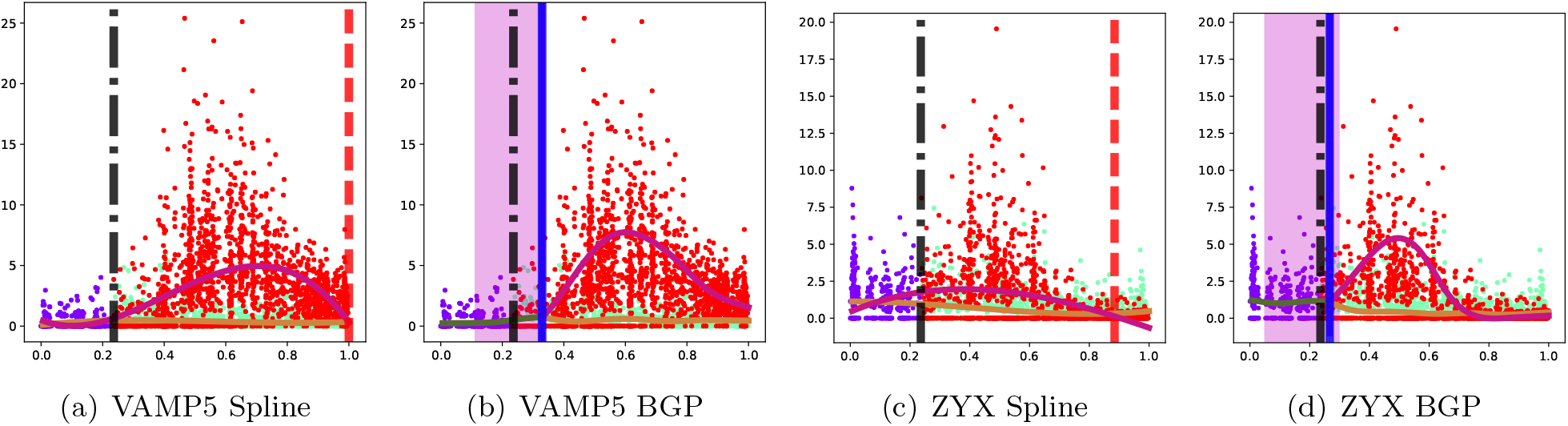
Hematopoiesis gene expression: contrasting spline and BGP fits under transitory gene expression. The global branching time (dashed black vertical line), spline branching point mode (red vertical line) and 98% posterior intervals (magenta vertical span) are also shown. The cells are marked according to the global allocation estimated by DDRTree.

## Conclusion

We have presented a flexible non-parametric probabilistic approach to robustly identify individual gene branching times. For scalability our model uses sparse variational inference implemented using the GPflow package. The probabilistic nature of our model allows for well-defined parameter estimation via maximisation of a bound on the marginal likelihood.

The spline model used by BEAM uses global branch assignments for each cell and is therefore unable to accurately identify branching times earlier than the global branching time. We found that branching time estimates from this spline-based approach were generally biased towards to global branching time. In contrast, the BGP method can robustly identify branching times as it estimates cell branch association for each gene independently while accounting for cell assignment uncertainty in the posterior branching times. We also found the BGP approach to be robust to global state estimation errors and high noise. The BGP branching time uncertainty can also be used in downstream analysis of the individual gene branching times; for example ranking genes in terms of their most likely or minimum branching times.

We have also included in our comparison a probabilistic linear method (Campbell and Yau, 2017). The linearity allows for an efficient joint estimation of both the pseudotime and global branching structure. Although this method does not estimate gene bifurcation times, a probabilistic estimate of an individual gene exhibiting branching behaviour is available. However, in our synthetic study we have found the pseudotime estimation not to be robust and this reduces the effectiveness of the method.

The application of the BGP method to the hematopoiesis data revealed the importance of modelling transitory gene expression which has the potential to confuse non-probabilistic methods. The model was able to automatically select the most likely branching location even in the presence of multiple crossing points in the gene expression without the need for any post-processing heuristics such as those included in the BEAM package.

Concurrent with our study, Penfold *et al*. have used changepoint kernels to develop similar branching Gaussian processes to identify bifurcations in single cell transcriptional datasets Penfold *et al*. (2017). They use a Markov chain Monte Carlo approach to estimate cell branch association and branching times. Their approach also explicitly models recombination, where individual branches are merged, and they can jointly estimate pseudotime. However, the computational complexity of their method would make application to genome-wide inference of branching times from unlabelled data challenging and that is the motivation for our sparse inducing point variational approach.

In the future we would like to extend our model to non-Gaussian likelihoods which would more accurately describe single cell data. This would increase inference complexity but could provide better calibrated uncertainty estimates. Another useful extension would be to jointly infer pseudotime and branching behaviour, which would also improve uncertainty estimation as the uncertainty arising from the estimation of the former would be included in the posterior branching uncertainty. Extending our model to multiple branching points is straightforward from a modelling standpoint but presents a more challenging optimisation problem wherein a tree prior on the branching structure may prove helpful (Simek *et al*., 2016). This extension would allow us to address the problem of selecting the correct number of branches in the global cellular branching dynamics.

## Methods

### Hematopoiesis RNA-seq data processing

The single-cell RNA-seq hematopoietic stem cell data was obtained from Paul *et al*. (2015). The data consists of 4423 cells. We applied the Monocle 2 algorithm (Qiu *et al*., 2017) and obtained the latent space depicted in Figure 5. Numerous minor branching events were identified by Monocle which is typical when no marker genes are used. We filtered out all cells assigned to the minor branching events resulting in a set of 3854 cells that were assigned to the main branching event. We removed all genes in which more than 80% of cells had zero count resulting in a set of 1343 genes that were examined in the branching analysis. Finally, to speed up computation we randomly selected a set of 900 cells and used *M* = 30 inducing points in our sparse BGP model.

### Branching model

We present in detail the probabilistic model description of the branching Gaussian process. We derive a lower bound on the model likelihood using variational inference techniques. Lastly we present a formulation of a sparse inducing point approximation that allows the application of the model to large datasets. We also discuss how to calculate the bound in a numerically stable manner and how to perform prediction on our model.

### Branching kernel

To model the branching process, we specify a branching kernel that constrains the latent branching functions to intersect at the branching point. We use a modified version of the kernel proposed in Yang *et al*. (2016) where the trunk *f* and branch kernel functions *g* and *h* are constrained to cross at the branching point *t_p_*. We place Gaussian process priors on all three functions and constrain them to intersect:

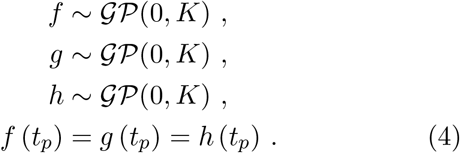

For simplicity the same kernel is used for all three functions although it would be straightforward to extend our framework to specify different kernels for each latent function. This would allow for instance, one branch to be modelled as a periodic function and the others as non-periodic. The extra flexibility would come at the cost of increasing the number of parameters that need to be estimated.

The resulting covariance between any two latent functions *f* and *g* constrained to cross at *t_p_* is

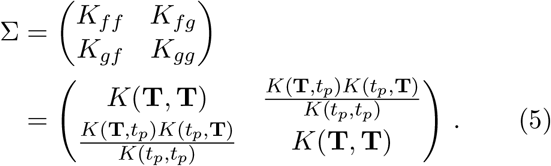

where *K*(**T**, **T**), *K*(**T**, *t_p_*) and *K*(*t_p_, t_p_*) are the kernel function evaluated between all training data pseudotimes **T**, between the training data and branching point, and solely at the branching point respectively.

In Yang *et al*. (2016) only two latent functions were specified, a control and perturbation condition where the former spanned the branching point. In our modified formulation three functions are used, allowing for discontinuity between the trunk and both branch latent functions. As an extension of our model, the derivatives of the latent functions could also be constrained to intersect at the branch point allowing for differentiable paths.

### Full GP inference

Let *Y* ∈ ℝ^*N*^ be the data of interest and let *M_f_* be the number of functions that are dependent. We specify a set of latent functions *F* for each data point of size *M ×* 1 where *M* = *N M_f_*^1^. Let *Z ∈* {0, 1*}^N ×M^* the binary indicator matrix which describes the association of each data point to a latent function. Each row or *Z* has only one non-zero entry. The model likelihood is

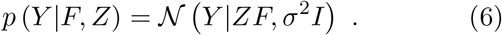

The extension to multiple independent outputs is straightforward as the likelihood factorizes

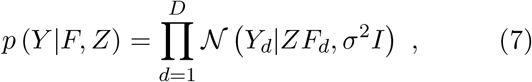

where *Y_d_* denotes the *N ×* 1 column vector of observations for output *d* and similarly *F_d_* denotes the *M ×* 1 column vector of latent function values. We omit the multiple output case from the derivation below for clarity.

As in Lázaro-Gredilla *et al*. (2012) we place a categorical prior on the indicator matrix *Z* and a GP prior on the latent functions *F*. Note that the latter does not factorize as in Lázaro-Gredilla *et al*. (2012) as we assume the latent functions are dependent:

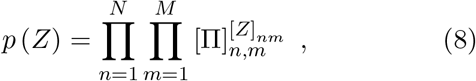

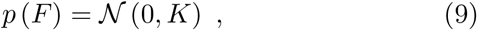

where for the multinomial distribution we have 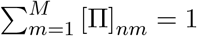 and *K* is the GP kernel^2^

The log likelihood is not analytically tractable as it involves integrating out the indicator matrix *Z*

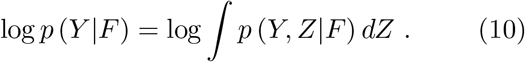

We proceed to compute a lower bound using Jensen’s inequality

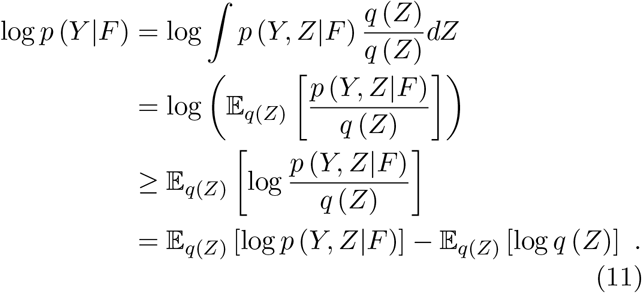

The last equation is usually presented in terms of a likelihood term and the tractable KL term

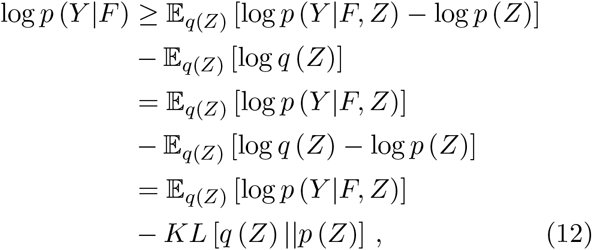

Where

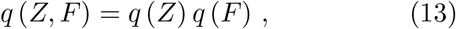

as by a mean-field assumption the latent functions *F* are independent of the association indicators *Z*. The log likelihood term is

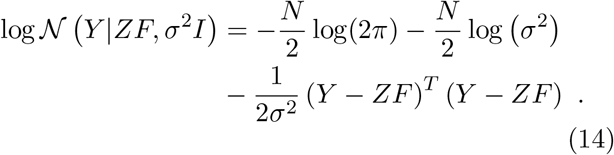

Taking the expectation with respect to the variational *n*=1 *m*=1distribution *q*(*Z*)

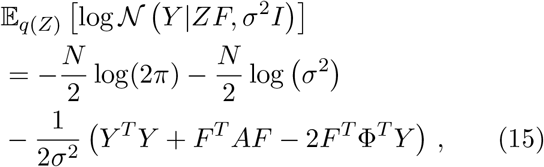

where we have defined

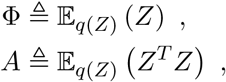

and the variational approximation is

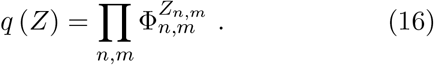

This encodes the mean-field assumption where we assume the posterior indicators factorize.

The second order expectation for *A* can be derived as follows: let *z_i_* the *N ×* 1 indicator vector for latent function *i* = *m*. We then have

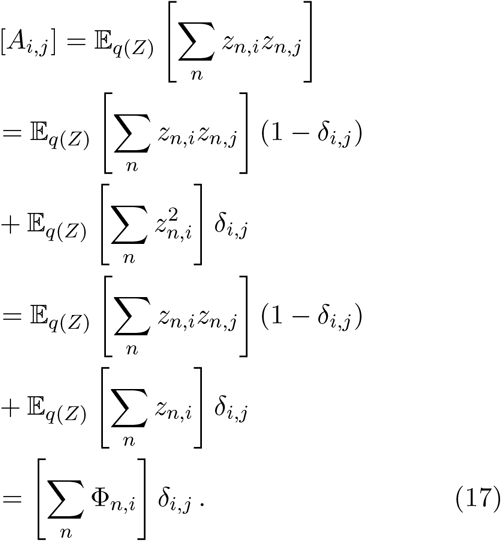

The third step follows from the fact the *z_n,i_* is binary and hence 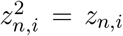 and we have used the notation *δ_i,j_* to denote the delta function which is 1 when *i* = *j* and 0 otherwise. The first term vanishes because cells cannot be assigned to more than one branch, and therefore *z_n,i_z_n,j_* = 0 when *i* ≠ *j*. In matrix notation the expectation is

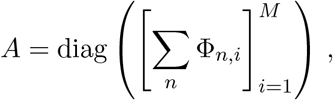

where diag denotes the diagonalisation of a vector and 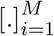 the construction of an *M* dimensional vector.

The KL divergence term is computable as

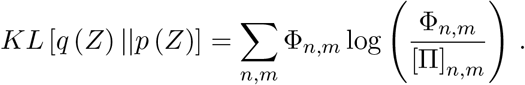

Our bound is therefore

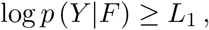

where we have defined

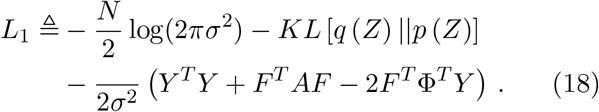

We proceed to integrate out the latent functions *F* to obtain the variational collapsed bound

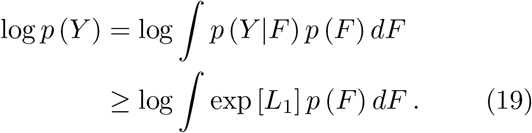

This bound holds because *L*_1_ is a bound to log *p* (*Y |F*) and the exponent function is monotonic. More details can be found in King and Lawrence (2006).

Setting the prior on the latent function as a GP log *p* (*F*) = log *N* (*F |*0, *K*) and substituting (18) into (19) results in the collapsed bound

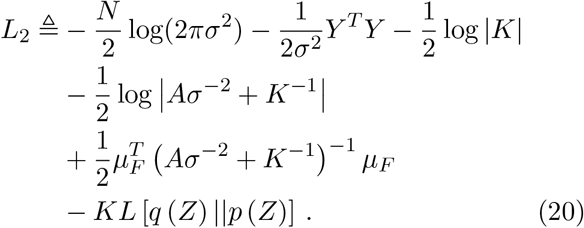

where we have *q* (*F*) = *N* (*µ_F_*, Σ_*F*_)

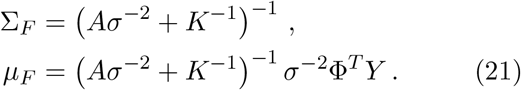

Expanding the quadratic term with the mean expression:

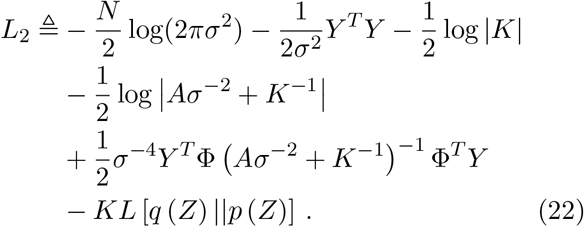

### Sparse GP inducing point approximation

To speed up inference we use a sparse inducing point approximation. This allows the algorithm to scale linearly with the number of training points. The number of inducing points is specified by the user and determines the trade-off between model accuracy and runtime; decreasing the number of inducing points will reduce runtime but will increase the approximation error.

The inducing points are treated as additional variational parameters which are optimised. The model specification is the same as before:

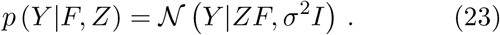

We introduce the inducing points *u* and explicitly specify their conditional relationship to the latent noise-free data *f*

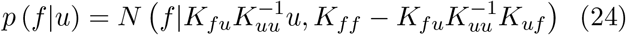

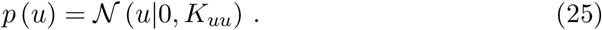

We now introduce the sparse GP approximation:

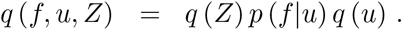

This is similar to the mean field approximation for the full model (Equation (13)) but we have introduced an additional mean-field factorisation *q* (*F*) = *p* (*f|u*) *q* (*u*).

The derivation proceeds as before with the resulting bound

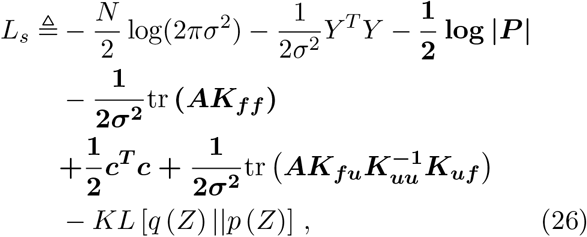

where the terms in bold are different compared to the full model bound (Equation (22)). In the bound above we have defined

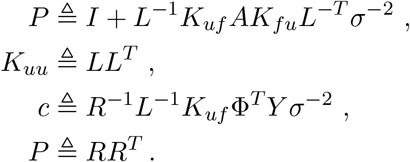

### Prediction

In the full model the predictive posterior a new point *f_∗_* is

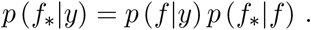

The training data posterior *p* (*f |y*) is approximated by *q* (*f*). To predict at a single *f_∗_* point we integrate over the latent functions values at the training data *f*,

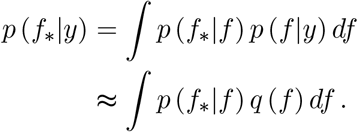

where 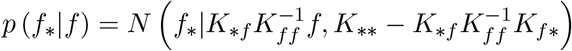 is the usual GP predictive density and *q* (*f*)=*N* (*f |µ_F_*, Σ_*F*_) as defined in Equation (21). The latter integral is tractable. Therefore there is no further approximation required past the original mean-field approximation (Equation (13)).

Similarly for the sparse approximation

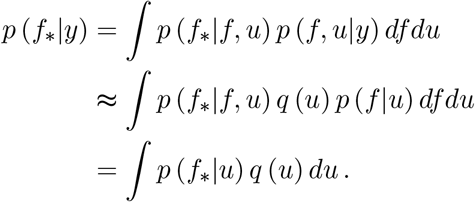

## Funding

MR and AB were supported by MRC award MR/M008908/1. JH was supported by an MRC fellowship.

## Availability of data and materials

An open source Python implementation of BGP is available at https://github.com/ManchesterBioinference/BranchedGP. Documentation and examples are available on the GitHub page. All data used in the paper are previously published and freely available.

## Competing interests

The authors declare that they have no competing interests.

## Author’s contributions

AB, JH and MR designed the algorithm. AB and JH implemented the code and AB performed the numerical experiments and analysis. AB and MR wrote the paper.

## Acknowledgements

We wish to thank Xiaojie Qiu for his help with the BEAM Monocle 2 code and Kieran Campbell with running the MFA model.

## Note

1 This expanded representation allows for efficient recomputation of the marginal likelihood for different branching times.

2 For simplicity we assume the same kernel for every output and latent trajectory function. Removing this restriction does not affect the derivation but will increase the inference complexity.

